# Estimating the heritability of psychological measures in the Human Connectome Project dataset

**DOI:** 10.1101/704023

**Authors:** Yanting Han, Ralph Adolphs

## Abstract

The Human Connectome Project (HCP) is a large structural and functional MRI dataset with a rich array of behavioral measures and extensive family structure. This makes it a valuable resource for investigating questions about individual differences, including questions about heritability. While its MRI data have been analyzed extensively in this regard, to our knowledge a comprehensive estimation of the heritability of the behavioral dataset has never been conducted. Using a set of behavioral measures of personality, emotion and cognition, we show that it is possible to re-identify the same individual across two testing times, and identify identical twins. Using machine-learning (univariate linear model, Ridge classifier and Random Forest model) we estimated the heritability of 37 behavioral measures and compared the results to those derived from twin correlations. Correlations between the standard heritability metric and each set of model weights ranged from 0.42 to 0.67, and questionnaire-based and task-based measures did not differ significantly in their heritability. We further derived nine latent factors from the 37 measures and repeated the heritability estimation; in this case, the correlations between the standard heritability and each set of model weights were lower, ranging from 0.15 to 0.38. One specific discrepancy arose for the general intelligence factor, which all models assigned high importance, but the standard heritability calculation did not. We present an alternative method for qualitatively estimating the heritability of the behavioral measures in the HCP as a resource for other investigators, and recommend the use of machine-learning models for estimating heritability.

## Introduction

Decades of research have accumulated abundant knowledge on the heritability of various human traits. A recent meta-analysis studied 28 functional domains and found the largest heritability estimates for several physical trait domains (such as the ophthalmologic and skeletal domains) but the lowest heritability for some psychological domains (such as the social values domain; Polderman et al., 2015). This domain-wise characterization was largely consistent with reported values from studies that focused on individual traits. For example, height is one of the most studied traits in the physical domain. An earlier study involving twins from eight countries estimated the heritability of height to be 0.87 - 0.93 for males and 0.68 - 0.84 for females (Silventoinen et al., 2003), although a more recent study of larger samples produced estimates up to 0.83 in boys and 0.76 in girls (Jelenkovic et al., 2016), comparable to the reported meta heritability of 0.73 (Polderman et al., 2015). By contrast, the heritability of psychological traits is generally estimated to be lower: episodic memory has a heritability around 0.3 – 0.6 (Papassotiropoulos and de Quervain, 2011) (with meta heritability around 0.6), and personality has a heritability around 0.4 (Vukasović and Bratko, 2015) (with meta heritability around 0.48). These traits have typically been studied in isolation in previous studies. Here we took advantage of the comprehensive set of measures available in the Human Connectome Project (HCP) dataset (including both self-report questionnaires and behavioral tasks), which allowed us to describe an individual’s psychological profile and similarity to others. Our goal was to apply modern machine learning methods to estimate heritability in this dataset, at the same time providing a resource that could be used for studies of heritability in the neuroimaging data component.

The Human Connectome Project (HCP) offers a uniquely rich sample of measures across the same 1200 subjects: structural, diffusion, and functional MRI, together with questionnaire- and task-based measures that assess many different psychological domains (Van Essen et al., 2013). The HCP dataset has proven to be a valuable resource for investigating individual differences. A number of recent studies have utilized the HCP dataset to predict personal identity, gender, fluid intelligence, personality, and executive function from brain connectivity (Dubois et al., 2018; Finn et al., 2015; Liu et al., 2018; Zhang et al., 2018).

Another valuable aspect of the HCP is that it has a rich and extensive family structure, including 149 genetically confirmed monozygotic twin pairs and 94 genetically confirmed dizygotic twin pairs. In principle, this provides a powerful resource for investigating the heritability of brain-behavior relationships. Several studies have used MRI data in the HCP to investigate the heritability of brain structures and connectivity patterns, many aspects of which are heritable (Ge et al., 2016). For instance, surface area and cortical thickness (Strike et al., 2019), the depth of Sulcal Pits (Le Guen et al., 2018), subcortical shape (Gutman et al., 2015), hippocampal subfield volumes (Patel et al., 2017) and cortical myelination (Liu et al., 2019) are all heritable structural features. Similarly, connectivity patterns, especially resting-state fMRI, have been shown to be heritable (Colclough et al., 2017; Adhikari et al., 2017), with highest estimates found for repeat measurements that account for transient fluctuations (Ge et al., 2017). Other studies have also probed the neural correlates of cognitive processes in the context of heritability using HCP data (Babajani-Feremi, 2017; Guen et al., 2018; Kochunov et al., 2016; Vainik et al., 2018). For instance, one study used bivariate genetic analyses to identify brain networks that were genetically correlated with cognitive tasks in math and language (Guen et al., 2018). Similarly, another study found common genetic influences for white matter microstructure and processing speed (Kochunov et al., 2016). Both studies demonstrated that heritability can provide a powerful link between brain and behavior.

Behavioral heritability is defined as the genetic contribution to the total variance for a phenotypic trait in a population, an important statistic for understanding individual differences. Twins (both monozygotic/MZ and dizygotic/DZ) are particularly useful for the estimation of heritability as they can help to differentiate the contribution of genes versus environment. In classical twin studies, the basic assumptions are that MZ twins share on average 100% of their alleles, while DZ twins share on average 50% of their alleles, and both MZ and DZ twins share a common environment. The total variance can be split into three components: additive genetics, shared environment and unique environment (often referred to as the ACE model) (Bouchard Jr and Propping, 1993; Falconer et al., 1996; Plomin et al., 1997). The simplest method for calculating heritability is to use Falconer’s formula. The formula assumes that unique environment contributes equally to the phenotypic variance for both MZ and DZ twins, and that therefore the difference between MZ phenotypic correlation and DZ phenotypic correlation arises solely because of genetic factors (Mayhew and Meyre, 2017; Polderman et al., 2015). Modern maximum likelihood-based modeling estimates various components for the total variance (Martin and Eaves, 1977; Winkler et al., 2015), but in essence relies on the same set of assumptions and logic, which are continually debated. The equal environment assumption (EEA), for example, is often believed to be violated. MZ twins, due to their physical resemblance, are likely to encounter a more similar social environment than DZ twins. Furthermore, gene-environment interaction is often not properly modeled or completely omitted as in the case of using Falconer’s formula in twin studies (Beckwith and Morris, 2008; Charney, 2017; Joseph, 2002; Kamin and Goldberger, 2002; Schönemann, 1997). Yet a recent meta-analysis paper that investigated the heritability of a wide range of human traits based on twin studies in the past fifty years showed that for 69% of the traits analyzed, there was a twofold difference in the MZ correlations relative to DZ correlations, consistent with a simple model that all twin resemblance was solely due to additive genetic variation (Polderman et al., 2015).

Given the lack of consensus on modeling the exact causes for the difference between MZ and DZ twins, we here present a model-free approach, using data-driven machine-learning tools. These have been shown to yield better results in the literature, most notably in improving the prediction of human phenotypic traits using single-nucleotide polymorphism (SNP) data (de Vlaming and Groenen, 2015; Koo et al., 2013; Mieth et al., 2016; Paré et al., 2017; Sun et al., 2008). One review that evaluated Ridge regression (which is a model used in our study) lists several advantages over conventional genome-wide association methods: (1) substantially increased accuracy, especially for large sample sizes; (2) the regularization term in the Ridge regression allows flexible accounting of the linkage disequilibrium between SNPs; (3) more computationally efficient than repeated simple regressions (de Vlaming and Groenen, 2015). Other models, such as Random Forest, a nonlinear machine learning model, have been used to predict coronary artery calcification using SNP data, achieving not only good prediction, but also reliably identifying best predictors across different datasets (Sun et al., 2008). Feature weights have been further utilized in one study that trained support vector machines (SVM) to classify siblings versus unrelated people using resting-state fMRI data to derive heritability for brain activity (Miranda-Dominguez et al., 2018). Overall, machine learning models have demonstrated superior prediction performance compared to conventional methods, and the feature weights learned by the models have the potential to be used for qualitative estimation of heritability.

The present study has two broad aims: 1, We tried to identify the same individuals and identical twins based on their behavioral profile, testing if the success in connectome fingerprinting that has been applied to the neuroimaging component of the HCP (Finn et al., 2015) could be replicated using this set of rich behavioral measures. 2, We set out to characterize the heritability of the behavioral data in this dataset using both the classical method and novel machine-learning based methods, for raw behavioral scores as well as nine latent factors. Aside from valuable comprehensive data on the heritability of psychological variables, our results can motivate hypotheses about the heritability of the neural underpinnings, which we hope future studies will pursue in the same subject sample.

## Materials and Methods

### Data

We used behavioral data from the Human Connectome Project (HCP) S1200 release under the domains of cognition, emotion and personality (Van Essen et al., 2013). The 37 selected variables were summary scores for either a behavioral task or a questionnaire (see Table S1 for more detailed description for each variable, and Figure 1A for their correlation structure). The NEO agreeableness score was re-calculated since item #59 was incorrectly coded at the time of downloading the data (an issue reported to and verified by HCP^1^). Since the variables were on different scales, we first pre-processed them to all have zero mean and unit variance. Each subject was thus essentially described by a vector of 37 scores/features, representing their psychological profile.

**Figure 1.**
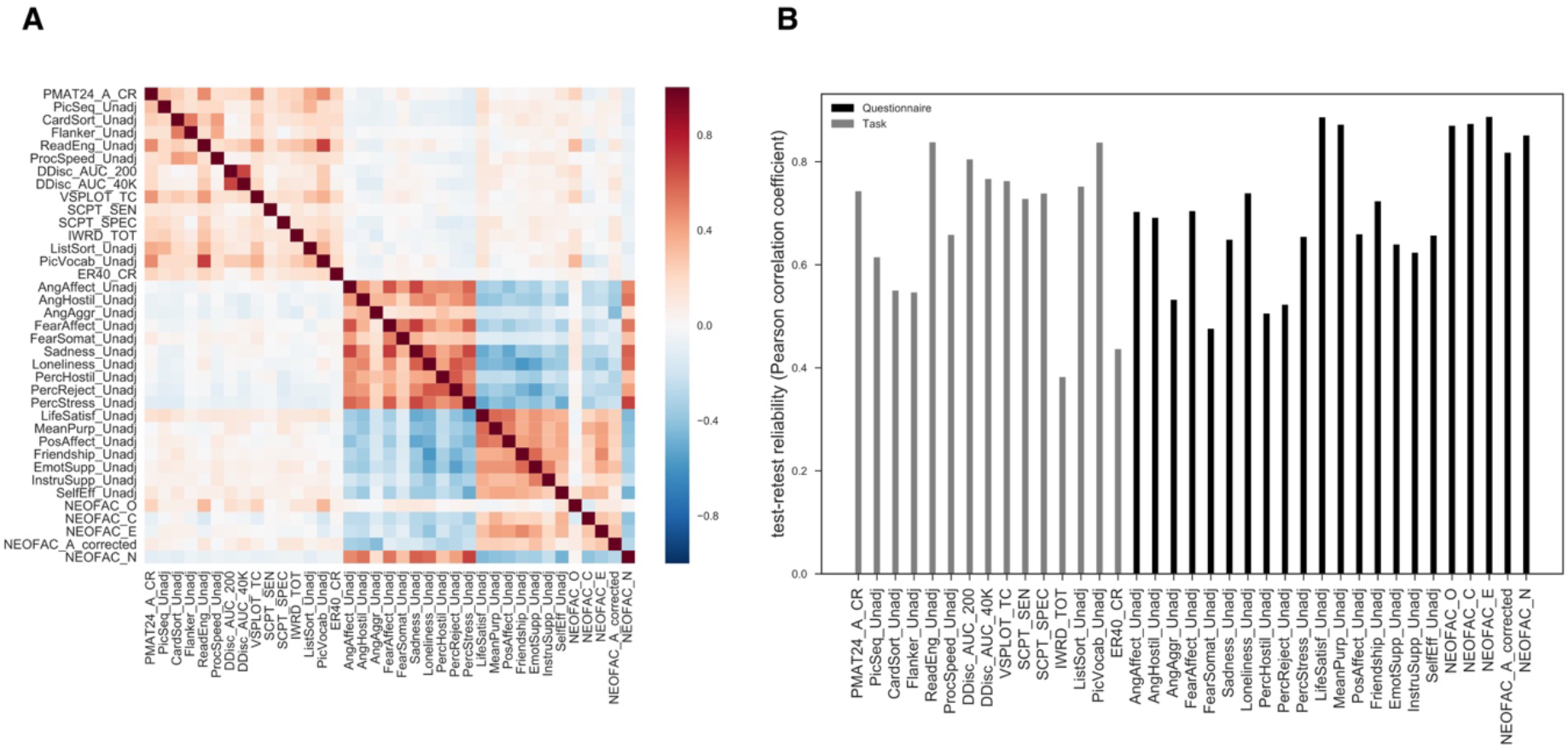
Overview of the dataset. (A) empirical correlation matrix for 37 behavioral variables in HCP (sample size N = 1189), color coded for Pearson’s correlation coefficient, (B) empirical test-retest reliability for 37 measures (sample size N = 46), color coded for domain. See inset legend for details. See Table S1 for descriptions of the variables.

Of 1206 subjects, 1189 subjects had complete data for the 37 scores of interest, and 1142 had family relationship data verified by genotyping, yielding a final set of 149 pairs of genetically confirmed monozygotic (MZ) twins (298 subjects, all of the same sex) and 90 pairs of dizygotic (DZ) twins (180 subjects, one twin pair was of opposite sex and thus excluded) with complete data for the 37 behavioral variables of interest. A subset of 46 MZ subjects had complete test-retest data for the selected 37 scores, which we used to calculate test-retest reliability (as their Pearson’s correlation coefficients, Figure 1B). We thus used 1189 subjects in total, of which 478 were either MZ or DZ twins.

### Same individual and twin identification

Same individual: We first asked how well a subject could be re-identified from their retest, compared to all other subjects, for the 46 subjects who had test-retest data available. We calculated pairwise Euclidean distances between a given subject’s retest data and each of the 1189 subjects’ original data (including the subject’s own original data) and then ranked the distances in ascending order to see if the subject’s retest data was closest to his/her own original data.

MZ twin: Similar to the above, we took one person (target) out of the 298 MZ twins and calculated pairwise Euclidean distances between this subject and each of the remaining 1188 subjects, and then ranked the distances in ascending order to see if the corresponding MZ twin was closest to the target.

### Standard calculation of heritability

In the behavioral genetics literature, a standard way to derive heritability is based on twin correlations calculated using Falconer’s formula (Falconer et al., 1996):

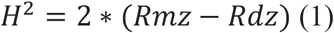

Where *H*^2^ is the overall heritability, *Rmz* the correlation for a phenotypic trait between monozygotic twins, and *Rdz* the correlation for a phenotypic trait between dizygotic twins.

### Machine learning approach

We took as input data the absolute feature-wise difference between each twin pair, described by a vector of 37 pre-processed behavioral variables as described above, giving us 149 MZ pair data and 90 DZ pair data which we tried to classify. To resolve unbalanced classes, we randomly sampled the DZ class with replacement to match the number of MZ cases.

We used three widely used models: a Ridge classifier, a simple univariate model, and a Random Forest model, which is a nonlinear decision tree-based model that ensures accurate feature weights even when features are correlated. For the univariate model, the dependent variable was the class and the independent variable was each of the 37 features; we used this simple model because it most clearly tests the maximal contribution of each feature in isolation.

We fitted both Ridge (the alpha parameter for the regularization term was determined by cross validation to be alpha = 100 for using 37 features, alpha = 10 for using the set of 9 factor scores calculated using linear regression, and alpha = 100 for using both sets of 18 factor scores) and Random Forest models (maximum tree depth was set to be 5 with 100 trees in the forest to prevent overfitting). Each model was estimated 1000 times; for each iteration, data was sampled as described above and then randomly split into 70% training data and 30% testing data. For Ridge classification, the testing accuracy and the coefficients for each of the 37 features were recorded. For Random Forest, the model returns feature importances that reflect mean decrease impurity (averaged across all decision trees in the random forest) (Leo et al., 1984). So, a feature with a higher importance score is better at decreasing node impurity (which is a metric of the number of mis-labeled data points at the current node of a decision tree), i.e., it is more informative than other features. We evaluated the performance of Random Forest models using both testing accuracy and ROC curve analysis.

### Factor analysis

Given the strong inter-correlations between the 37 behavioral variables (Figure 1A) and the consideration that a single individual variable/task will yield an imprecise measure of the underlying psychological construct, we performed an exploratory factor analysis using SPSS with principal axis factoring as the extraction method, and kept nine factors that had eigenvalues >1, which together explained about 60% of the variance. Factors were rotated using Promax rotation, since there was no evidence that the factors were orthogonal. We also calculated the factor scores using both regression and Bartlett methods.

### Statistical testing

The statistical significance of our identification tests was evaluated with permutation testing. Over 1000 iterations, subject identity was randomly shuffled from the original dataset across the 1189 subjects, and the same identification procedures described above (both same-individual identification and identical-twin identification) were performed to derive the empirical distribution for chance-level identification accuracy.

To assess the statistical significance of our classification performance, we constructed the 95% confidence interval from the empirical testing accuracy distribution (resulting from the 1,000 bootstraps that we performed) for each classification problem. A bootstrap p-value was also computed as the ratio of the instances of having a testing accuracy equal or lower than 50% (which is the expected chance accuracy for random guessing with equal probability for a balanced binary classification) out of the total number of bootstraps.

Permutation testing was also used to test for a significant difference in average heritability between the questionnaire domain and behavioral task domain. The null hypothesis was that the task and the questionnaire domain comprised the same distribution. Under the null hypothesis, the number of all possible permutations (selecting 15 out of 37 measures as task scores) was 9.4*10^2^, which we approximated using Monte Carlo sampling of 100,000 permutations. For each permutation, we randomly assigned 15 values to the task domain and the rest to the questionnaire domain and then calculated the absolute difference between the two heritability means as our test statistic. Statistical significance was quantified as the probability (under the null hypothesis) of observing a value of the test statistic more extreme than what was actually observed. We performed the same analysis for four sets of heritability estimates (heritability calculated using Falconer’s formula, univariate model weights, Ridge weights, and feature importances for the Random Forest model, each consisting of 37 values). For heritability calculated using Falconer’s formula, we set any negative value to be zero.

## Results

### Same individual and Monozygotic twin identification based on psychological profiles

Given the rich behavioral measures, we first attempted to re-identify the same individual using all of the 37 measures. Of the 46 subjects with retest data, we were able to re-identify 26, yielding an accuracy of 56.5 % with a median distance rank of 1.0 and a mean distance rank of 12.1 among 1189 people. We performed permutation testing to assess the statistical significance of our identification accuracy. Across 1,000 iterations, the highest success rate achieved was 2/46 which is roughly 4.3% and the p-value associated with obtaining at least 26 correct identifications was <0.0001.

We carried out the same analysis for MZ twin identification: compared to other siblings and genetically unrelated people, MZ twins should be most similar to one another (Bouchard Jr and Propping, 1993; Falconer et al., 1996; Plomin et al., 1997). Of the 298 MZ subjects, we identified the exact corresponding MZ twin for 21 of them, yielding an accuracy of 7.0 % with a median distance rank of 47.5 among 1188 people. Assessing statistical significance with 1000 permutations, the highest success rate achieved was 3/298, roughly 1.0%, and the p-value associated with obtaining at least 21 correct identifications was <0.0001. Thus, our ability to identify somebody’s identical twin based on the behavioral data was considerably worse than our ability re-identify the same individual (7% accuracy vs. 56.5%), even though statistically highly significant.

The ability to re-identify a given individual from test-retest essentially sets an upper bound on our ability to identify a MZ twin, and presumably reflects the specific limitations of this particular dataset, including factors such as the number of features (37 compared to ideally infinite) and the reliability of the features (test-retest reliability in Figure 1B). We next investigated the heritability of each measure and the fundamental assumptions in twin studies.

### The standard method of calculating heritability

In twin studies, the most common approach to calculate heritability is to compare the difference in correlations between MZ and DZ twins (see Introduction). In this framework, we calculated the heritability using Falconer’s formula (Figure 2A). As can be seen from the figure, the heritability calculated in this manner had a very large range across the different tasks and actually yielded a negative value for two of them (MZ correlation was smaller than the DZ correlation). This demonstrates some of the flaws with using Falconer’s formula on this dataset. One possible explanation for this theoretically invalid result could be that the measures have poor test-retest reliability. Yet, for the two tasks in question, the short Penn line orientation test had a test-retest reliability of 0.76 and the life satisfaction questionnaire had a test-retest reliability of 0.89. Another limiting factor could be the sample size used to calculate the twin correlations (on the order of 100 here). There exist more complex modeling approaches to estimate heritability (Martin and Eaves, 1977; Winkler et al., 2015), but fundamentally, those methods rely on the same assumptions. Given the patent limitations of the standard approach, which is well known in the literature (Beckwith and Morris, 2008; Charney, 2017; Joseph, 2002; Kamin and Goldberger, 2002; Mayhew and Meyre, 2017; Schönemann, 1997), we took an alternative approach of estimating heritability, which is to make use of machine learning models that are more data-driven and less model-based.

**Figure 2.**
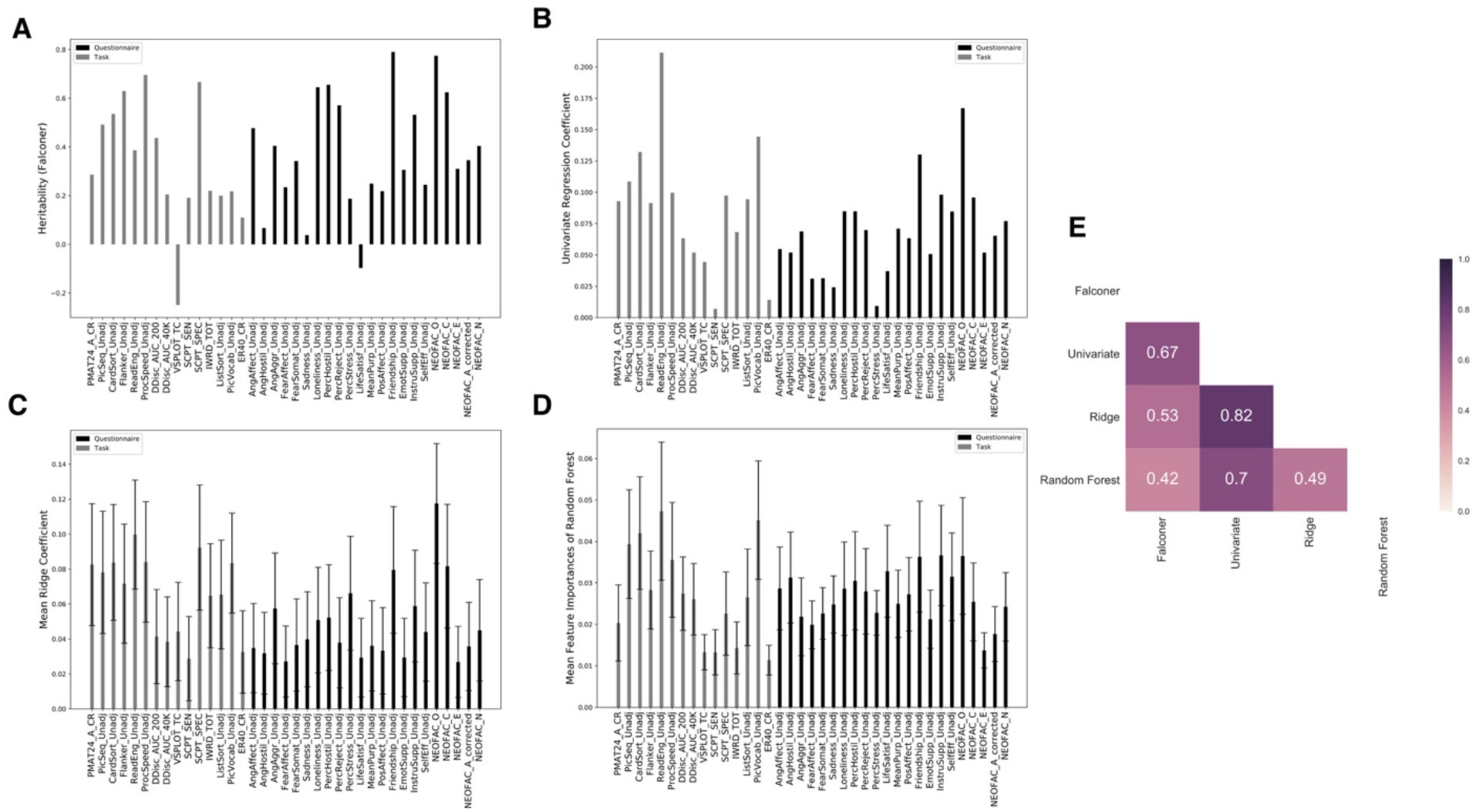
Heritability estimation across four methods for 37 behavioral measures. (A) heritability calculated using Falconer’s formula (note that for VSPLOT and LifeSatisf, Rmz is smaller than Rdz and thus negative heritability.); (B) univariate coefficients for each feature; (C) mean feature coefficients averaged across 1000 iterations for Ridge classifier (error bars represent standard deviation of coefficients); (D) mean feature importances averaged across 1000 iterations for Random Forest (error bars represent standard deviation of importances); (E) correlation matrix for four sets of heritability estimates assigned to 37 measures, color coded for Spearman’s rank correlation. See inset legend for details.

### A machine learning alternative for estimating heritability for the 37 measures

The traditional approach derives heritability from the differences between MZ and DZ twins. If we assume that any differences between the two types of twin pairs indeed arise solely from genetics, then a classifier trained to distinguish MZ twins and DZ twins should assign greater weights to the features that have higher heritability, as they are more informative for discriminating the two classes. This allows us to test at least qualitatively how reasonable the heritability estimations were that we derived above using standard methods.

The first approach we used was Ridge classification, which is a variant of a simple multivariate model with a regularization term that forces the weights to be more stable and robust to correlated features (Freckleton, 2011; Gopakumar et al., 2016) (which was the case for the measures we selected as illustrated in Figure 1A). The mean coefficients for each feature are plotted in Figure 2C, the model had a mean testing accuracy of 68.7% (95% confidence interval for the testing accuracy: [58.9%,77.8%]; the bootstrap p-value under the null hypothesis that testing accuracy is not significantly higher than 50% was <0.0001). In addition to Ridge regression, we also fitted the simplest univariate model for each of the 37 measures, an OLS regression model with a single feature, each one of the coefficients are shown in Figure 2B. This univariate regression would therefore reflect the maximal contribution from each feature in isolation, allowing a clearer quantification of each individual feature’s heritability than the Ridge or Random Forest models, which incorporate multicollinearity between features. The two sets of coefficients (univariate and Ridge) had a Spearman’s rank-order correlation of 0.82 across the 37 features.

Another popular approach is the Random Forest classifier, which is a nonlinear model comprised of many decision trees. For each decision tree inside the forest, the method draws a randomly sampled training set and only considers a random sample of features for splitting at each node. The structure of the model helps with the problem of highly correlated features and allows more stable and accurate estimations of feature weights (importances). The mean feature importances are plotted in Figure 2D, the model had a mean predictive accuracy of 79.4% (95% confidence interval: [71.1%,87.8%]; p <0.0001); mean area under the ROC curve was 0.88 (with a standard deviation of 0.04).

To compare all these different results, we quantified the correlations between all four sets of values, including classic heritability as calculated from Falconer’s formula, Ridge classifier coefficients, univariate model coefficients and Random Forest feature importances. We found good agreement across different approaches with Spearman’s rank correlation ranging from 0.42 to 0.82 (Figure 2E), demonstrating the validity of our novel machine-learning approach for estimating heritability qualitatively. Considering that we had correlated features in the dataset (Figure 1A), the results also partially confirmed the capability of both Ridge and Random Forest at handling feature correlations as they both agreed well with the univariate coefficients, correlated at 0.82 and 0.7 respectively. Results that corrected for test-retest reliability were similar to the uncorrected ones presented here (Figure S1).

We next asked a more general question: are the heritability or feature weights on average significantly different for the behavioral task domain compared to the self-report questionnaire domain? Under the null hypothesis that average heritability for the task and the questionnaire domain are not significantly different, we constructed the distribution of the absolute difference for average heritability between the task and questionnaire domain (Figure 3), and calculated the p-values for four sets of heritability estimates (see more details in the method section). For all cases except Ridge (for which the p-value was 0.021, uncorrected for testing our hypothesis with the four sets of heritability estimates), we found no strong evidence to reject the null hypothesis. When taking test-retest reliability into consideration by simple disattenuation (dividing by rest-retest reliability), again only Ridge coefficients had the smallest p-value of 0.008 (Figure S2). However, it may not be valid simply to divide by test-retest reliability, since measures with very poor reliability could yield artificially inflated heritability. As noted above, a single task or questionnaire is often limited in reflecting the meaningful psychological variable of which it is a measure, and we therefore next conducted factor analysis to derive latent factors across our 37 measures.

**Figure 3.**
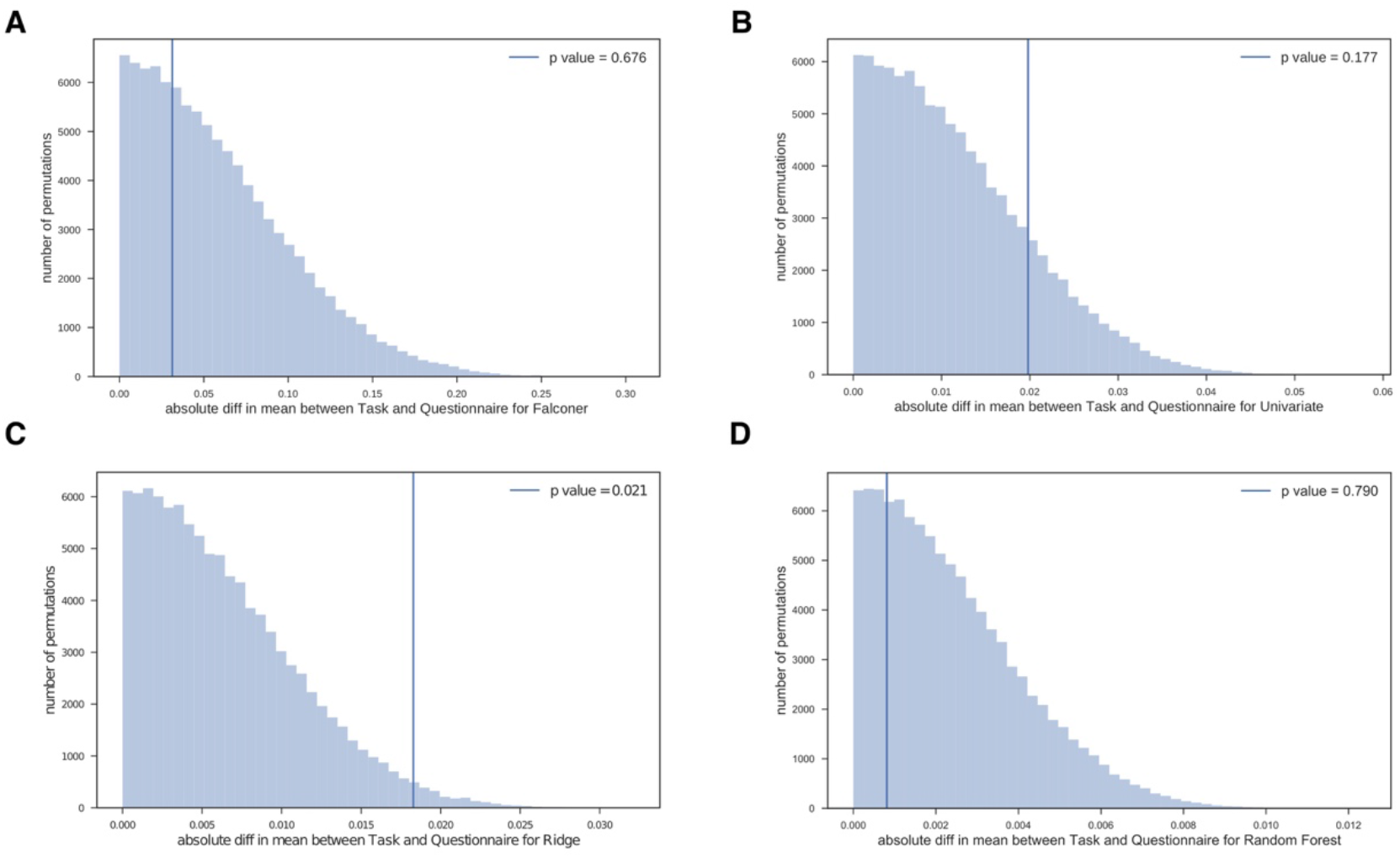
Distribution of the absolute mean difference between the task and questionnaire domain (vertical line indicates actual observation of the difference for average heritability between the task and questionnaire domain) for (A) heritability calculated using Falconer’s formula; (B) univariate coefficients for each feature; (C) Ridge classifier coefficients; (D) Random Forest feature importances.

### Estimating heritability for the factors

We extracted nine factors from all 37 measures that together accounted for 59.7% of the total variance (Table S2). The interpretations and accounted variances of the factors were factor 1: positive social ability (22.2%); factor 2: negative affect (11.0%); factor 3: general intelligence (5.1%); factor 4: self-regulation (4.7%); factor 5: attention and processing speed (4.0%); factor 6: agreeableness (3.6%); factor 7: self-efficacy (3.2%): factor 8: language and communication (3.2%) and fac9: competitiveness (2.8%).

We also computed factor scores using both regression and Bartlett methods for reliability (since factor scores are indeterminate). These two methods produced two sets of very similar factor scores for the same nine factors (see correlation structure between 18 factor scores in Figure S3). We used these two set of factor scores simultaneously as features in the Ridge classifier and Random Forest model to further assess the ability of each model to handle highly correlated features (a more challenging task than handling the 37 variables which were less inter-correlated in comparison). For a model that’s robust to correlation among features, it should be able to assign similar weights or importances to features that are highly correlated to each other.

We repeated the previous analyses using both sets of factor scores so that each subject was represented by a vector of 18 factor scores to derive standard heritability, Ridge coefficients, univariate coefficients and Random Forest feature importances for the nine factors (Figure S4). For Heritability using Falconer’s formula and univariate coefficients (Figure S4 A,B), each factor score was treated independently, so they were not susceptible to the influence of correlation among factors. For the Ridge classifier, for the two sets of factor scores, the two sets of coefficients (Figure S4 C) had a Pearson’s correlation of 0.79. For the Random Forest analysis, the correlation between the two sets of feature importances (Figure S4D) was 0.61. Therefore, these results further confirmed that Ridge and Random Forest were able to assign similar weights to highly correlated features and that their estimation of heritability was reliable.

We repeated the analysis for the Ridge classifier and Random Forest using only the one set of factor scores derived from regression methods (Figure 4C, D). When using the nine regression factor scores alone, The Ridge classifier had a mean accuracy of 64.2% (95% CI: [55.3%,73.3%]; bootstrap p-value = 0.006) while the Random Forest classifier had a mean testing accuracy of 77.9% (95% CI: [67.8%,86.7%]; bootstrap p-value <0.0001) and mean area under the ROC curve of 0.86 (with a standard deviation of 0.04). The reduction of model performance compared to using all 37 measures was minimal, indicating that the latent factors captured the information relevant to estimating heritability. For the set of factor scores derived by regression, when trained alone versus together with the other set of factor scores computed by the Bartlett method, the Spearman’s rank correlation of Ridge coefficients was 0.73. For the Random Forest classifier, the feature importances were correlated at 0.93. These results demonstrated that the feature weights that Ridge and Random Forest learned for the nine factors (calculated using Regression method) were robust and consistent.

**Figure 4.**
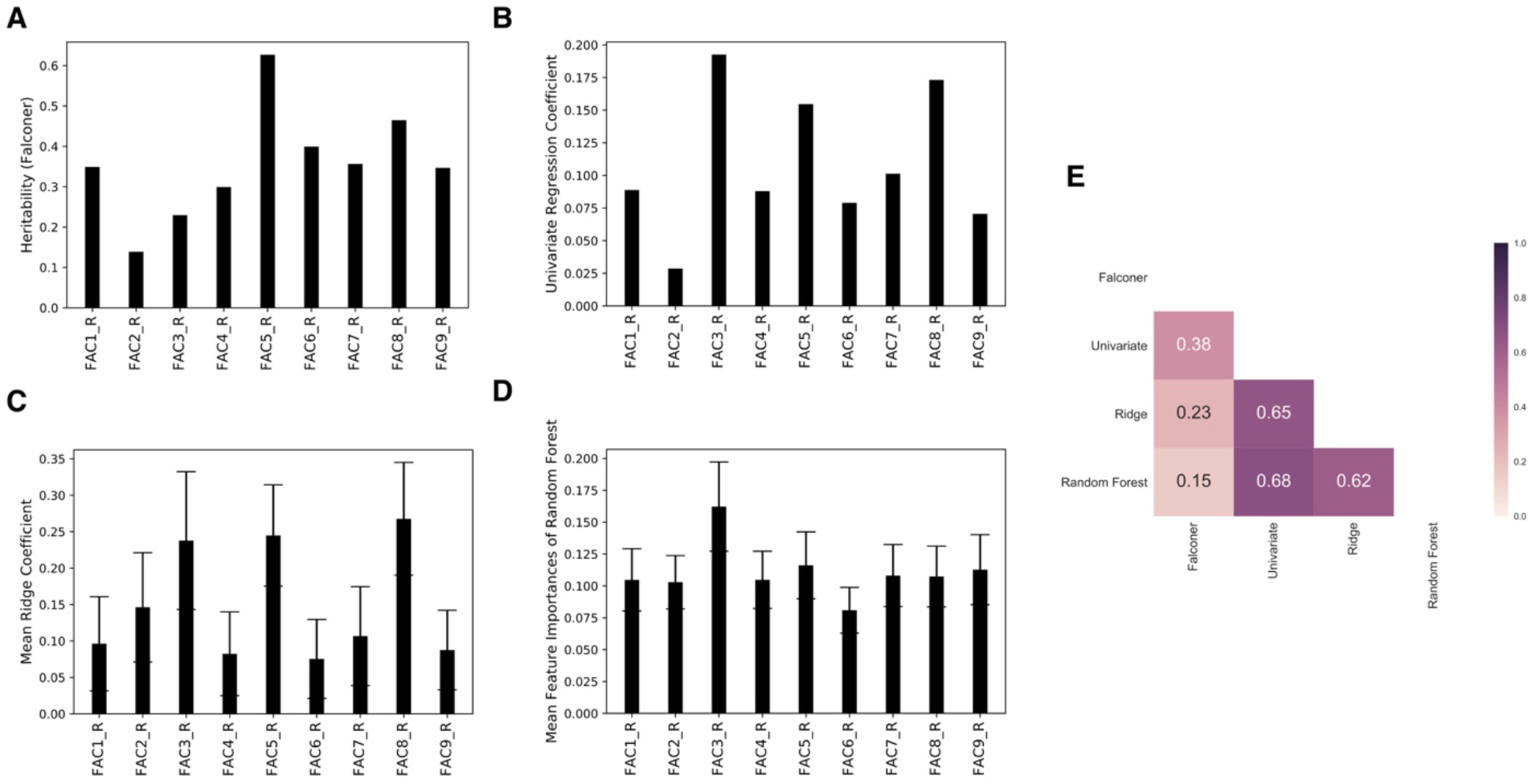
Heritability estimation across four methods for nine latent factors. (A) heritability calculated using Falconer’s formula; (B) univariate coefficients for each factor; (C) mean feature coefficients averaged across 1000 iterations for Ridge classifier (error bars represent standard deviation of coefficients); (D) mean feature importances averaged across 1000 iterations for Random Forest (error bars represent standard deviation of importances); (E) correlation matrix for four sets of values assigned to 9 factors, color coded for Spearman’s rank correlation.

Recall that for the 37 measures, standard heritability and feature importances from the three models agreed relatively well, from 0.42 to 0.67 (Figure 2E). However, for the nine factors, the classical heritability estimates from Falconer’s formula (Figure 4A) had lower correlations with the three other sets of model estimation, from 0.15 to 0.38 (Figure 4E). One specific difference, for example, was the estimation of factor 3 which reflects general intelligence. All three models assigned high importance to this factor while the traditional heritability calculation assigned a rather low value at 22.9%. In the literature, the estimation for the heritability of intelligence is quite high, often above 50% and sometimes reported to be as high as 80% (Bouchard, 2004; Panizzon et al., 2014; Plomin and Deary, 2015). The machine-learning models are thus likely to have produced a more accurate estimation of heritability from this dataset than the standard formula was able to.

## Discussion

### Summary of results

In this study, we analyzed a comprehensive set of 37 behavioral scores in the Human Connectome Project. When representing each subject using this set of behavioral data, we were able to achieve a behavioral fingerprinting accuracy of 56.5% for individuals, and in the case of identifying identical twins, an accuracy of 7.0% (both significantly above chance). We further computed heritability for those 37 scores in two general schemes: classical correlation-based method using Falconer’s formula, and three machine-learning based methods (univariate linear model; Ridge classify and Random Forest model), and found relatively high correlations between the two schemes (Figure 2E). Given the inter-correlations among the 37 scores, an exploratory factor analysis was conducted to extract nine latent factors, whose heritability we assessed similarly. In this case, the correlations between the classical method and machine-learning-based ones were lower (Figure 4E).

### Individual and MZ twin identification

Our behavioral fingerprinting scheme was inspired by the success of connectome fingerprinting using HCP data (Finn et al., 2015). Our accuracy of 56.5% was relatively high considering the limiting factors that we faced: a small number of features compared to the connectome fingerprinting (which had 268 nodes and 35778 edges) and measurement error from some measures with relatively low test-retest reliability. Our identification of MZ twins faced the same limitations, but we observed a drop of performance to an accuracy of 7.0%. This accuracy drop alone would seem to put a limit on the strength of the heritability of our measures.

One possible explanation is that the unique environment actually accounts for a substantial portion of the variance for those measures, overwhelming the contribution of common environment and genes. According to a study that used maximum likelihood modeling, unique environment does account for the majority of variances for many of the measures in the HCP, including some of the ones we selected (Winkler et al., 2015). This may also partly explain the modest classification accuracy of Ridge classification between MZ twins and DZ twins, since stronger contribution of unique environment implies weaker contribution of genetics and common environment to the overall phenotypic variances, thus diminishing group differences between MZ twin pairs and DZ twin pairs.

### Comparison of the standard correlation-based method versus machine-learning based methods of estimating heritability

The standard analysis calculates the heritability based on the difference between MZ and DZ correlations for a phenotypic trait. One immediate shortcoming of this approach is that it can sometimes yield negative heritability in cases where the MZ correlation is actually smaller than the DZ correlation. In our case, we found that two measures that had good test-retest reliability had negative heritability using Falconer’s formula. Possible reasons for negative heritability could be due to small sample size and/or lack of explicit knowledge of the common environment. However, it should be mentioned that a negative estimation of heritability is not rare using such methods and although most researchers attribute such invalid results to noise, they could in fact be evidence against the assumptions behind the calculations (Schönemann, 1997; Steinsaltz et al., 2018).

We therefore developed an alternative approach to estimate heritability, that is, to train machine learning models to distinguish MZ twin pairs and DZ twin pairs. If the ACE model stands, then measures/features that have high heritability would be assigned larger weights since they are more informative for the classification. We found good rank correlations between the standard heritability and another three sets of model coefficients for the 37 behavioral variables (Figure 2E). However, when applied to nine latent factors, the agreement between the standard heritability and another three sets of model coefficients were substantially lower (Figure 4E). However, the three machine learning models had good agreement with one another, as shown by relatively high rank correlations (all above 0.6) (Figure 4E). As mentioned above, the standard heritability estimation for the general intelligence factor deviated greatly from the other three models, and from the literature. Such disagreement raises concerns about the validity of the assumptions made by the ACE model and the usage of traditional methods for calculating heritability, leading us to recommend the use of machine learning methods to estimate heritability empirically.

### Limitations and future directions

To the best of our knowledge, this is the first application of utilizing machine learning models to estimate heritability for behavioral measures using the HCP data. We will evaluate each model respectively and make recommendations for future usages.

For the univariate linear model, a conceptually simple model, each measure was evaluated independently for its maximal contribution for the classification. For both raw measures and latent factors, univariate model coefficients agreed best with standard heritability calculations. Though it should be noted that given the shortcomings of standard calculations that we discussed before, good agreement with these doesn’t necessarily imply agreement with the true set of heritability values.

The second model we used was a Ridge classifier, a commonly used linear model to deal with correlated features (Dormann et al., 2013; Freckleton, 2011; Gopakumar et al., 2016). A recent paper (using single-nucleotide polymorphism data) concludes that Ridge classification will improve predictive accuracy substantially compared to standard repeated univariate regression for a large enough sample size (de Vlaming and Groenen, 2015). In our case, we also wanted to derive accurate coefficients, as estimation of heritability. As a regularized regression, Ridge has proven to be effective at handling feature correlation, illustrated by its good agreement with the univariate coefficients (Figure 2E, Figure 4E) and its ability to assign similar weights to the two sets of factor scores (Figure S4 C).

The Random Forest model was also robust with respect to correlations among features (e.g., Figure S4 D, for two sets of almost identical factor scores for the same nine factors, the two sets of feature importances had a Pearson’s correlation of 0.61), and achieved the highest accuracy for the classification between MZ twin pairs and DZ twin pairs. Given the nonlinear nature of the model, though, the feature importances should be interpreted in a qualitative sense rather than in an absolute sense.

In this study, we focused on the classification of MZ twins versus DZ twins as a starting point, because within the standard ACE framework, the model weights in this classification scheme have a clear theoretical interpretation (that they should only reflect heritability). Within the assumptions of the ACE model, weights derived from classification of MZ twins versus genetically unrelated people, for example, would reflect a complex mixture of genetic effects and common environment, which would be difficult to interpret. However, future research could explicitly quantify the common environment (the HCP does not provide such information, besides household ID), and even propose new models to explain the composition of the total phenotypic variance. Researchers could then train multiple classifiers (such as MZ versus DZ, full siblings versus half siblings) to further disambiguate the contribution of each component.

This general machine-learning framework could be applied to the heritability estimation of brain activation as well, a source of data much more mined in the HCP than the phenotypic data. One recent study organized a subset of HCP subjects into MZ twins, DZ twins, siblings and unrelated people and found greater activation pattern similarity with greater genetic relatedness (Etzel et al., 2019). Using our approach, such findings could go beyond simple association to heritability estimation, by training classifiers on brain activation patterns for different groups. In summary, the machine learning methods that we introduced here have the potential to not only supplement standard heritability calculations, but also to provide insights for theories explaining phenotypic variance, and studies that focus on linking brain activation with behavior.

## Conflict of interest

The authors declare that the research was conducted in the absence of any commercial or financial relationships that could be construed as a potential conflict of interest.

## Author contributions

Y.H. and R.A. developed the overall general analysis framework. Y.H. conducted all final analyses and produced all figures. Y.H. and R.A. wrote the manuscript.

## Funding

Funded by NSF grant BCS-1840756 and BCS-1845958.

## Acknowledgements

Data were provided [in part] by the Human Connectome Project, WU-Minn Consortium (Principal Investigators: David Van Essen and Kamil Ugurbil; 1U54MH091657) funded by the 16 NIH Institutes and Centers that support the NIH Blueprint for Neuroscience Research; and by the McDonnell Center for Systems Neuroscience at Washington University.

## Footnotes

1 https://www.mail-archive.com/hcp-users@humanconnectome.org/msg06007.html

