## supplemental figures and tables for "Estimating the heritability of psychological measures in the Human Connectome Project dataset"

### *Supplementary Material*

#### Supplementary Figures and Tables

##### Supplementary Figures

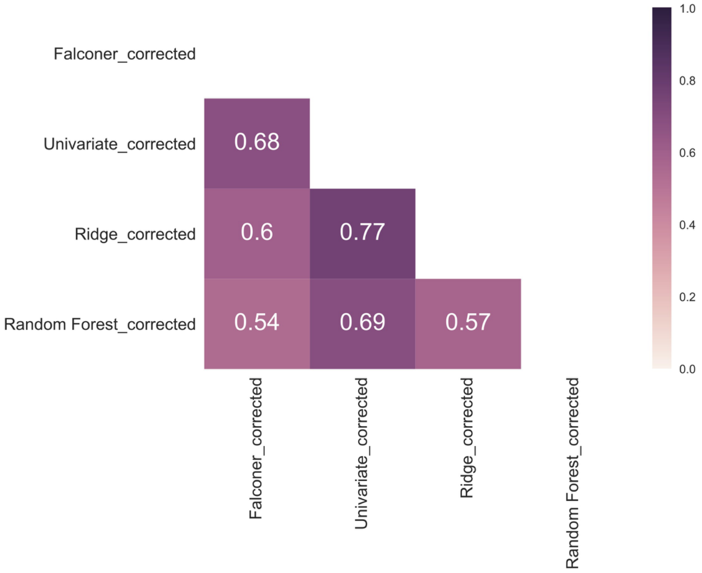


**Figure S1.** Spearman's rank correlation matrix for four sets of heritability estimates assigned to 37 measures that are corrected for test-retest reliability.

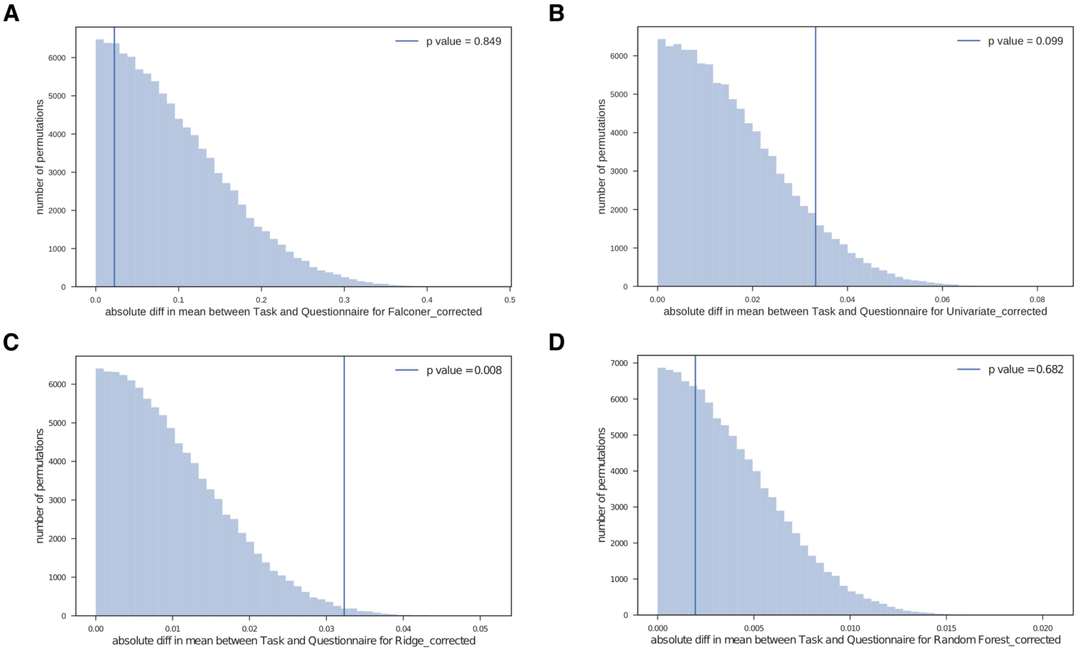


**Figure S2.** Distribution of the absolute mean difference between the task and questionnaire domain (vertical line indicates actual observation) for (A) heritability calculated using Falconer's formula; (B) univariate coefficients for each feature; (C) Ridge classifier coefficients; (D) Random Forest feature importances. All values are corrected for test-retest reliability.

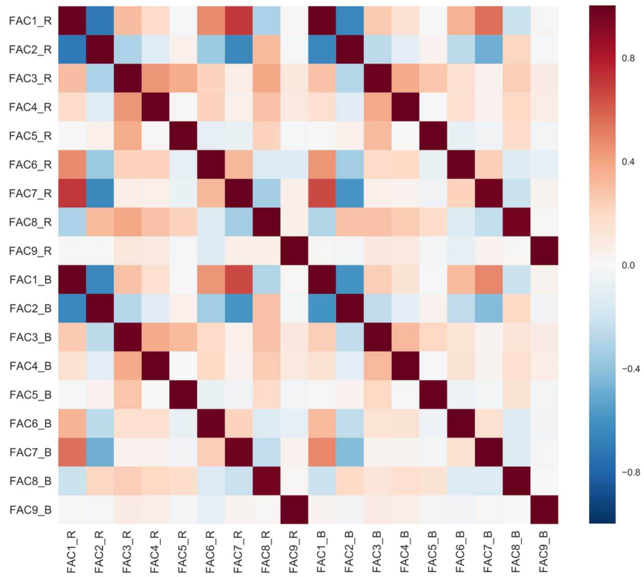


**Figure S3.** Pearson's correlation matrix for two sets of factor scores derived using regression method (FAC1_R to FAC9_R) and Bartlett method (FAC1_B to FAC9_B).

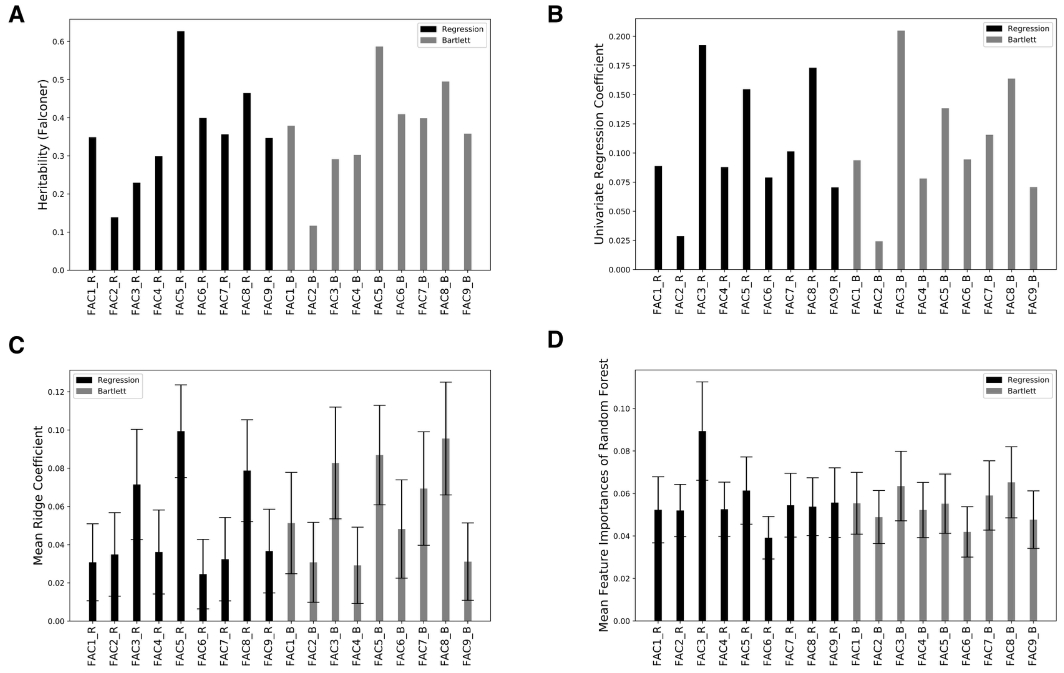


**Figure S4.** (A) heritability calculated using Falconer's formula; (B) univariate coefficients for each feature; (C) mean feature coefficients averaged across 1000 iterations for Ridge classifier (error bars represent standard deviation of coefficients); (D) mean feature importances averaged across 1000 iterations for Random Forest (error bars represent standard deviation of importances for two sets of factor scores (color coded for Regression and Bartlett).

#### Supplementary Tables

**Table S1.** List of 37 behavioral variables selected and their basic descriptions.

| HCP variable name | Description | Task/ questionnaire |
|---|---|---|
| PMAT24_A_CR | Penn Progressive Matrices | Task |
| PicSeq_Unadj | NIH Toolbox Picture Sequence Memory Test | Task |
| CardSort_Unadj | NIH Toolbox Dimensional Change Card Sort Test | Task |
| Flanker_Unadj | NIH Toolbox Flanker Inhibitory Control and Attention Test | Task |
| ReadEng_Unadj | NIH Toolbox Oral Reading Recognition Test | Task |
| ProcSpeed_Unadj | NIH Toolbox Pattern Comparison Processing Speed Test | Task |
| DDisc_AUC_200 | Delay Discounting: $200 | Task |
| DDisc_AUC_40K | Delay Discounting: $40,000 | Task |
| VSPLOT_TC | Variable Short Penn Line Orientation | Task |
| SCPT_SEN | Short Penn Continuous Performance Test: Sensitivity | Task |
| SCPT_SPEC | Short Penn Continuous Performance Test: Specificity | Task |
| IWRD_TOT | Penn Word Memory Test | Task |
| ListSort_Unadj | NIH Toolbox List Sorting Working Memory Test | Task |
| PicVocab_Unadj | NIH Toolbox Picture Vocabulary Test | Task |
| ER40_CR | Penn Emotion Recognition Test | Task |
| AngAffect_Unadj | NIH Toolbox Anger-Affect Survey | Questionnaire |
| AngHostil_Unadj | NIH Toolbox Anger-Hostility Survey | Questionnaire |
| AngAggr_Unadj | NIH Toolbox Anger-Physical Aggression Survey | Questionnaire |

| | | |
|---|---|---|
| FearAffect_Unadj | NIH Toolbox Fear-Affect Survey | Questionnaire |
| FearSomat_Unadj | NIH Toolbox Fear-Somatic Arousal Survey | Questionnaire |
| Sadness_Unadj | NIH Toolbox Sadness Survey | Questionnaire |
| Loneliness_Unadj | NIH Toolbox Loneliness Survey | Questionnaire |
| PercHostil_Unadj | NIH Toolbox Perceived Hostility Survey | Questionnaire |
| PercReject_Unadj | NIH Toolbox Perceived Rejection Survey | Questionnaire |
| PercStress_Unadj | NIH Toolbox Perceived Stress Survey | Questionnaire |
| LifeSatisf_Unadj | NIH Toolbox General Life Satisfaction Survey | Questionnaire |
| MeanPurp_Unadj | NIH Toolbox Meaning and Purpose Survey | Questionnaire |
| PosAffect_Unadj | NIH Toolbox Positive Affect Survey | Questionnaire |
| Friendship_Unadj | NIH Toolbox Friendship Survey | Questionnaire |
| EmotSupp_Unadj | NIH Toolbox Emotional Support Survey | Questionnaire |
| InstruSupp_Unadj | NIH Toolbox Instrumental Support Survey | Questionnaire |
| SelfEff_Unadj | NIH Toolbox Self-Efficacy Survey | Questionnaire |
| NEOFAC_O | NEO-FFI Openness to Experience | Questionnaire |
| NEOFAC_C | NEO-FFI Conscientiousness | Questionnaire |
| NEOFAC_E | NEO-FFI Extraversion | Questionnaire |
| NEOFAC_A | NEO-FFI Agreeableness | Questionnaire |
| NEOFAC_N | NEO-FFI Neuroticism | Questionnaire |

**Table S2.** Pattern matrix for factor analysis. The Table shows the loadings of each of the 37 measures (rows) onto the 9 factors that we derived.

| Pattern Matrix | | | | | | | | | |
|---|---|---|---|---|---|---|---|---|---|
| | Factor | | | | | | | | |
| | 1 | 2 | 3 | 4 | 5 | 6 | 7 | 8 | 9 |
| NEOFAC_O | 0.110 | 0.040 | 0.073 | 0.001 | -0.042 | 0.052 | 0.024 | 0.543 | 0.018 |
| NEOFAC_C | -0.087 | -0.006 | -0.010 | -0.012 | 0.003 | 0.121 | 0.562 | -0.141 | -0.013 |
| NEOFAC_E | 0.525 | 0.152 | -0.236 | -0.030 | 0.094 | 0.093 | 0.322 | 0.226 | 0.102 |
| NEOFAC_A_corrected | 0.038 | 0.020 | 0.026 | -0.037 | -0.029 | 0.735 | 0.114 | 0.128 | -0.028 |
| NEOFAC_N | 0.061 | 0.445 | -0.136 | 0.003 | 0.007 | -0.003 | -0.495 | -0.044 | -0.040 |
| PMAT24_A_CR | -0.082 | 0.061 | 0.668 | 0.004 | -0.020 | -0.020 | 0.045 | 0.119 | -0.001 |
| PicSeq_Unadj | 0.017 | 0.105 | 0.525 | -0.049 | 0.091 | -0.017 | 0.008 | -0.188 | 0.002 |
| CardSort_Unadj | -0.023 | -0.004 | 0.192 | -0.010 | 0.676 | 0.021 | 0.107 | -0.051 | -0.037 |
| Flanker_Unadj | 0.033 | -0.106 | -0.056 | 0.040 | 0.721 | 0.023 | -0.062 | 0.050 | 0.057 |
| ReadEng_Unadj | -0.020 | -0.039 | 0.570 | 0.003 | -0.018 | 0.079 | -0.131 | 0.383 | 0.032 |
| ProcSpeed_Unadj | 0.114 | 0.048 | 0.176 | 0.017 | 0.502 | -0.054 | -0.046 | -0.150 | 0.005 |
| DDisc_AUC_200 | -0.016 | 0.005 | -0.073 | 0.809 | -0.002 | -0.046 | 0.020 | 0.041 | 0.041 |
| DDisc_AUC_40K | -0.031 | 0.045 | -0.045 | 0.890 | 0.043 | 0.035 | -0.018 | -0.032 | -0.069 |
| VSPLOT_TC | -0.049 | 0.090 | 0.462 | 0.033 | 0.103 | 0.006 | 0.080 | 0.173 | -0.030 |
| SCPT_SEN | 0.033 | 0.066 | 0.212 | -0.039 | 0.058 | -0.046 | 0.026 | -0.024 | 0.017 |
| SCPT_SPEC | 0.000 | 0.161 | 0.454 | 0.005 | -0.087 | 0.056 | 0.043 | -0.131 | -0.083 |
| IWRD_TOT | 0.008 | 0.026 | 0.366 | -0.030 | 0.040 | 0.133 | -0.094 | -0.016 | 0.016 |
| ListSort_Unadj | -0.087 | 0.003 | 0.555 | -0.040 | 0.026 | -0.104 | 0.063 | 0.016 | -0.017 |
| PicVocab_Unadj | 0.081 | -0.129 | 0.503 | 0.042 | -0.041 | 0.010 | -0.276 | 0.402 | 0.043 |
| ER40_CR | -0.152 | 0.064 | 0.371 | -0.077 | 0.085 | 0.080 | 0.109 | -0.016 | -0.057 |
| AngAffect_Unadj | 0.053 | 0.826 | 0.136 | 0.003 | -0.060 | -0.224 | 0.069 | -0.013 | 0.070 |
| AngHostil_Unadj | -0.113 | 0.247 | -0.079 | -0.046 | 0.063 | -0.213 | -0.249 | -0.027 | 0.070 |
| AngAggr_Unadj | 0.012 | 0.091 | -0.034 | -0.024 | -0.035 | -0.555 | 0.030 | 0.021 | 0.044 |
| FearAffect_Unadj | 0.093 | 0.964 | 0.105 | 0.011 | -0.031 | 0.128 | 0.018 | -0.003 | 0.033 |
| FearSomat_Unadj | 0.167 | 0.626 | 0.158 | 0.028 | -0.006 | -0.095 | 0.016 | -0.036 | 0.098 |
| Sadness_Unadj | -0.186 | 0.794 | 0.061 | 0.043 | -0.011 | 0.078 | 0.068 | 0.071 | -0.082 |

| | | | | | | | | | |
|---|---|---|---|---|---|---|---|---|---|
| Loneliness_Unadj | -0.582 | 0.170 | 0.015 | 0.040 | 0.013 | 0.067 | -0.133 | 0.051 | 0.146 |
| PercHostil_Unadj | -0.203 | 0.248 | -0.062 | -0.035 | 0.025 | -0.167 | -0.014 | 0.088 | 0.409 |
| PercReject_Unadj | -0.570 | 0.170 | -0.129 | -0.026 | 0.044 | 0.024 | 0.053 | 0.094 | 0.430 |
| PercStress_Unadj | -0.092 | 0.550 | -0.173 | -0.043 | 0.034 | 0.075 | -0.229 | 0.076 | 0.054 |
| LifeSatisf_Unadj | 0.532 | -0.082 | 0.216 | 0.029 | -0.022 | 0.034 | -0.049 | -0.175 | 0.321 |
| MeanPurp_Unadj | 0.477 | 0.099 | 0.027 | 0.062 | -0.071 | 0.101 | 0.231 | -0.124 | 0.279 |
| PosAffect_Unadj | 0.585 | -0.166 | -0.112 | -0.030 | 0.040 | 0.037 | -0.009 | 0.038 | 0.272 |
| Friendship_Unadj | 0.835 | 0.094 | -0.147 | -0.037 | 0.062 | -0.049 | 0.067 | 0.199 | -0.147 |
| EmotSupp_Unadj | 0.938 | 0.126 | -0.033 | -0.017 | 0.005 | 0.046 | -0.133 | 0.067 | -0.104 |
| InstruSupp_Unadj | 0.682 | 0.066 | 0.007 | 0.006 | 0.019 | -0.048 | -0.149 | -0.026 | -0.018 |
| SelfEff_Unadj | 0.223 | -0.111 | 0.074 | 0.031 | -0.037 | -0.200 | 0.507 | 0.223 | 0.015 |
